# Intercellular contact and cargo transfer between Müller glia and to microglia precede apoptotic cell clearance in the developing retina

**DOI:** 10.1101/2023.10.06.561302

**Authors:** Michael Morales, Anna P Findley, Diana M. Mitchell

## Abstract

To clarify our understanding of glial phagocytosis in retinal development, we used real-time imaging of larval zebrafish to provide cell-type specific resolution of this process. We show that radial Müller glia frequently participate in microglial phagocytosis while also completing a subset of phagocytic events. Müller glia (MG) actively engage with dying cells through initial target cell contact and phagocytic cup formation after which an exchange of the dying cell from MG to microglia often takes place. Additionally, we find evidence that Müller glia cellular material, possibly from the initial Müller cell’s phagocytic cup, is internalized into microglial compartments. Previously undescribed Müller cell behaviors were seen, including cargo splitting, wrestling for targets, lateral passing of cargo to neighbors, and engulfment of what is possibly synaptic puncta. Collectively, our work provides new insight into glial functions and intercellular interactions, which will allow future work to understand these behaviors on a molecular level.

**Summary Statement:** Real-time imaging of developing zebrafish retinas reveals intercellular exchanges between Müller glial cells and to microglia during the clearance of apoptotic cells.

## Introduction

Cell death occurs in tissues and organs during development and throughout the life of an animal and these dying cells require clearance by other cells through phagocytosis. The process of phagocytosis has been well-studied in a variety of cells and tissues, and in vertebrates there is a class of professional phagocytes known as macrophages which, as tissue residents, efficiently perform this function across the body (Gordon and Plüddemann, 2017). Within the central nervous system, the specialized population of resident macrophages known as microglia perform phagocytosis, as well as additional specialized functions, in this very complex and delicate neural tissue. Recently, the function of microglia has risen to greater interest in the contexts of homeostasis, pathology, and regeneration. The phagocytic function of microglia is indeed well-appreciated, yet also in the central nervous system exist neuroglial cells that have become known to engage in phagocytic activity (Jung and Chung, 2018). Within the retina in particular are the Müller glia, which have been reported to engage in phagocytosis in a variety of species and contexts (Bailey et al., 2010; Lew et al., 2022; Nomura-Komoike et al., 2020; Sakami et al., 2019; Thiel et al., 2022b). Despite this body of work, the extent of such phagocytic activity is not yet fully defined and has not been directly observed in vivo in real-time.

The retina is the neural component of the eye and serves to convert incoming visual stimuli into the experience of vision through the use of specialized neurons. Müller glia are neuroglial cells that extend radially across the retina, spanning the full breadth of the retina’s neuronal components (Güngör Kobat and Turgut, 2020). Müller cells are known to crucially support the neurons that they neighbor by providing various homeostatic services such as metabolite and neurotransmitter synthesis and recycling (Pfeiffer et al., 2020). Zebrafish Müller cells have also garnered a special interest due to their ability to produce neuronal progenitors leading to regeneration of the retina following damage in zebrafish (Bernardos et al., 2007; Fausett and Goldman, 2006; Nagashima et al., 2013; Thummel et al., 2008). Further, Müller glia-microglia interactions appear to influence such regenerative potential (Fischer et al., 2014; Fogerty et al., 2022; Todd et al., 2020; White et al., 2017). Phagocytosis or engulfment of dying neurons by retinal Müller glia has been described in several studies through static visualization of dying cell markers within Müller cells (Bailey et al., 2010; Egensperger et al., 1996; Krylov et al., 2023; Lew et al., 2022; Morris et al., 2005; Nomura-Komoike et al., 2020; Sakami et al., 2019; Thiel et al., 2022b). In some cases, these reports seem to contrast with evidence that microglia dominate clearance of dying cells during retinal development (Blume et al., 2020; Francisco-Morcillo et al., 2014; Thiel et al., 2022b) or following induced retinal damage (Mitchell et al., 2018; White et al., 2017). Our recent work demonstrated the ability of Müller glia to substantially increase phagocytosis in the absence of microglia in an otherwise normally developing retina (Thiel et al., 2022b). However, to our knowledge to date, studies reporting Müller glia phagocytosis in vivo involve primarily fixed tissue or static samples and the full process of Müller glial phagocytosis in vivo, in real-time, has not been documented.

When differentiated, Müller glial bodies are stationary within the inner nuclear layer of the retina. Because Müller glial processes enwrap neurons, there remains the possibility that engulfment occurs as Müller processes collapse around already enwrapped neurons as they die. Alternatively, Müller glia may actively sense and reach dying cells with cellular processes that then enclose around the target. In addition, given that we and others reported that microglia are the dominant phagocyte in the developing CNS (Blume et al., 2020; Casano et al., 2016; Mazaheri et al., 2014; Möller et al., 2022), there remains a strong possibility that interactions between microglia and other cell types such as the Müller glia occur during, and to facilitate, the clearance process. While early endosomal markers support active engulfment of dying cells by Müller glia in the absence of microglia (Thiel et al., 2022b), lysosomal fusion of such compartments upon engulfment within Müller glia has not been clearly demonstrated. In summary, static images do not have the power to sufficiently reveal Müller glia phagocytic propensity or to visualize transient interactions of Müller glia with microglia and/or dying cells.

We used timelapse in vivo imaging of transgenic zebrafish reporter lines to address these outstanding questions by simultaneously recording Müller glia, microglia, and dying cells in the developing retina. Through these recordings, we demonstrate that Müller glia sense, contact, and enwrap dying cells via cellular processes extended from the Müller cell body. We find that most clearance events involve intercellular interactions between Müller glia and microglia with the same dying cell, whereby Müller glia most frequently make the first contact via phagocytic cup formation, after which the target is then transferred to microglia for terminal engulfment and degradation. We detected both membrane-localized and cytoplasmic fluorescent Müller glial reporters inside of presumptive microglial phagocytic compartments, suggesting that along with the dying cell cargo, cellular material from Müller glia is transferred to microglia, possibly through the cargo transfer/engulfment process. Completion of engulfment of dying cells by the Müller glia is also evident in real-time for a subset of phagocytic events. We also observed numerous behaviors of Müller glia that have not been previously described, including engulfment of phosphatidyl-serine (PtdSer) positive puncta in the presumptive inner plexiform layer. In situ hybridization revealed that microglia strongly express numerous PtdSer receptors while Müller glial expression is limited and heterogenous. Our real-time observations reconcile seemingly contradictory reports of Müller glia vs microglia phagocytosis of dying cells, suggest that clearance of dying cells in the vertebrate retina involves both Müller glia and microglia, and reveal previously undocumented, remarkably dynamic behaviors and intercellular interactions of Müller glia with microglia and their neighbors that should initiate an evolution of our view of how these glial cells function.

## Results

### Müller glia actively and dynamically extend processes towards dying cells

To visualize Müller glia, microglia, and dying cells simultaneously during retinal development, we performed time-lapse imaging of triple transgenic larval zebrafish expressing fluorescent protein reporters for Müller glia (*TP1:mTurquoise*), microglia (*mpeg1:mCherry* (Ellett et al., 2011)), and apoptotic cells (*TBP:Gal4; UAS:SecA5-YFP*, henceforth referred to as “SecA5- YFP” (Blume et al., 2020; van Ham et al., 2010)). The *TP1:mTurquoise* line reports Notch signaling via the TP1 transcriptional element, which labels Müller glia in the developing retina after ∼48 hours post fertilization (hpf) due to sustained Notch signaling activity in Müller glia (MacDonald et al., 2015). The SecA5-YFP reporter marks exposed phosphatidyl serine (PtdSer) on the surface of presumptively apoptotic cells by using YFP fused to AnnexinA5 (AnnA5). Exposure of PtdSer is an early apoptotic event, (Martin et al., 1995) thus the SecA5-YFP reporter allows visualization of apoptotic cell bodies via YFP (Herzog et al., 2019; Mazaheri et al., 2014; van Ham et al., 2010; van Ham et al., 2012). Zebrafish microglia are visualized using *mpeg1* driven transgenes (Blume et al., 2020; Ellett et al., 2011; Mitchell et al., 2018). Zebrafish larvae were imaged beginning at an age of ∼2.5 days post fertilization (dpf), or approximately 58 hours post fertilization (hpf) for a total duration of 10 hours, at 3-minute intervals, coincidental with a wave of apoptosis in the retina previously described (Biehlmaier et al., 2001), and which we previously visualized with the *mpeg1* and SecA5-YFP reporters (Blume et al., 2020).

Consistent with our previous study (Blume et al., 2020), microglia were seen to engulf YFP+ cell bodies throughout the retina (Movie 1). In addition, now observable due to the Müller glia reporter, we observed that Müller glia (MG) were active and dynamic, extending processes from the MG cell body towards SecA5-YFP+ targets in their vicinity (Movie 1, Movie 2, Figure 1). These processes extended around the body of a YFP+ cell, often forming a compartment-like structure resembling a phagocytic cup (Figure 1A, 1B, 1C). Process extensions originating from Müller glia directed towards a dying cell primarily originate in the region of the Müller cell closest to the dying cell, rather than a more distally derived extension (Figure 1 A-C). The extension of processes from Müller glia towards and around dying cells was not confined to any particular region of the retina. We observed the contact of MG processes with dying cells in both the apical, inner, and basal regions of the retina, suggesting that the Müller glia respond to death of numerous types of neurons (Figure 1 A-C). Consistent with death of inner retinal cells during this time window (Biehlmaier et al., 2001), we observe cell death arising in regions consistent with ganglion cells, amacrine cells, and bipolar cells (inner retina). Notably, however, we did not observe SecA5-YFP+ Müller glia in any of our recordings, nor did we observe compaction of any Müller glia process or cell bodies, indicating that Müller glia themselves do not undergo a wave of cell death during this timeframe.

**Figure 1:**
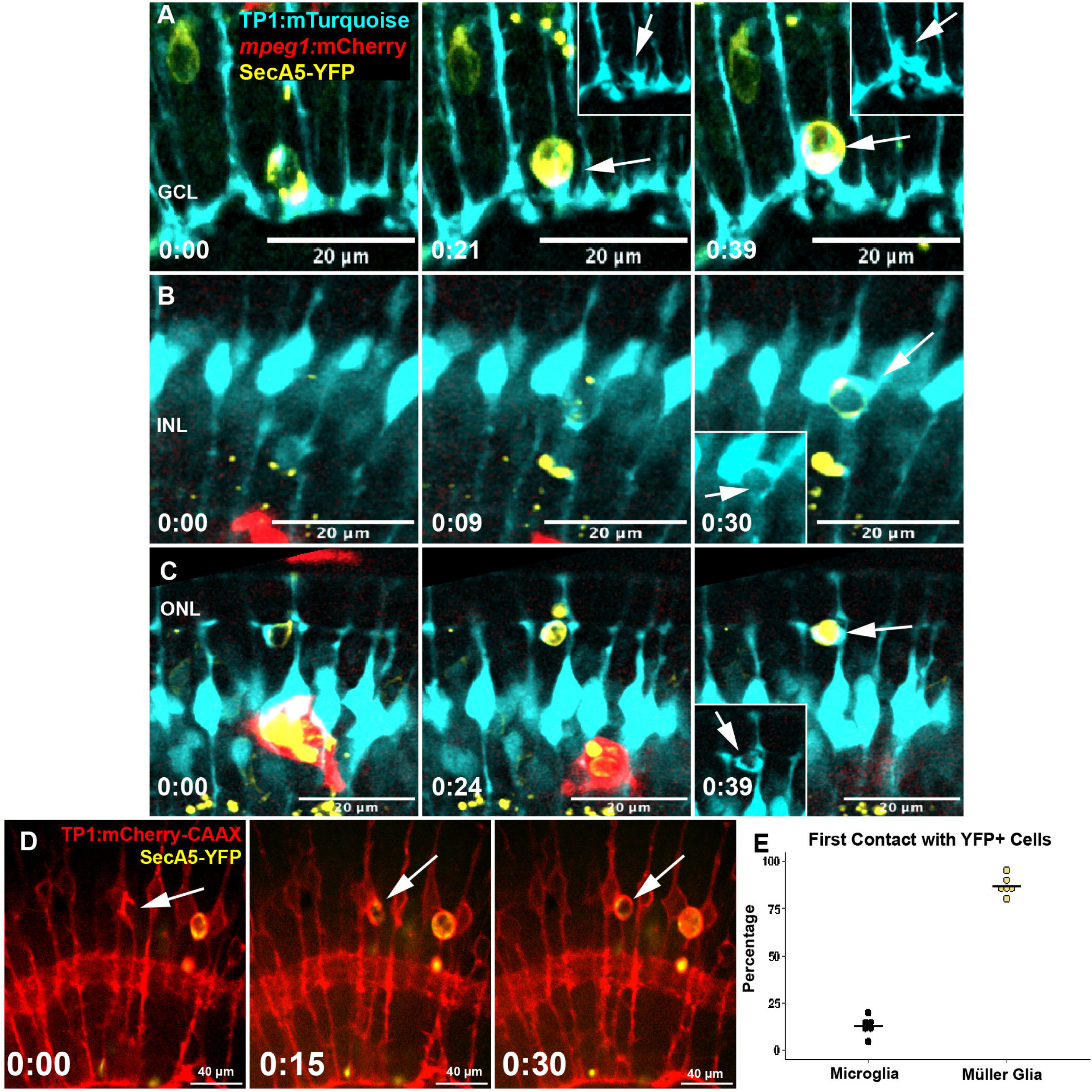
Müller cell process extension and contact with dying cells. A: Time-lapse stills show a Müller cell (*TP1:mTurquoise*) recognizing a dying cell in the ganglion cell layer (SecA5-YFP+) and extending a process towards and around the emerging YFP+ cell. Insets at 0:21 and 0:39 (hours:minutes) shows Müller cells forming phagocytic cup-like structures around the YFP+ cells. B: Time-lapse stills show MG process extension around YFP+ cell in the inner nuclear layer. Inset at 0:30 shows phagocytic cup structure (arrow). C: Time-lapse stills show an extension of MG processes around a YFP+ cell in the outer retina. Inset at 0:39 shows the phagocytic cup structure (arrow). D: Membrane reporter expressed by Müller glia (TP1:mCherry-CAAX) further demonstrates the extension of processes around SecA5-YFP+ dying cells forming a phagocytic cup, often enwrapping the YFP+ cell (arrows). In this example, the extension of the mCherry+ cellular processes begins prior to (timestamp 0:00) the appearance of the YFP signal (timestamp 0:15). Timestamp, bottom right (hr:min). E: Quantification of percentages of YFP+ apoptotic cell bodies initially contacted by microglia versus Müller cells in each time-lapse recording. Timestamps, bottom left (hr:min), relative to the first frame in the time series.

Because the *TP1:mTurquoise* transgene encodes a cytoplasmic reporter expressed in Müller glia, we wanted to visualize a fuller extent of this behavior using a membrane-tagged reporter. To do this, we used the reporter line *TP1:mCherry-CAAX* (Thiel et al., 2022b), in which Müller glia express a membrane-tagged mCherry reporter. We coupled the *TP1:mCherry-CAAX* reporter with the SecA5-YFP reporter. In a parallel manner, with this reporter system, we again visualized Müller cell processes extending towards and around the YFP+ dying cell (Figure 1D). Interestingly, process extension from the Müller cell often initiated prior to detection of the YFP+ cell death reporter (Figure 1D), suggesting that the Müller glia respond to signals released from the dying cell as cell death is executed. Given the observed response of Müller glia to dying cells, and the known role of microglia in clearance during this time (Blume et al., 2020), we quantified the first cell type initially reaching to and engaging SecA5-YFP+ cell bodies in our set of recordings from *TP1:mTurquiose*;SecA5-YFP;*mpeg1:mCherry* triple transgenic fish. This analysis revealed that the majority (>75%) of phagocytic events occurring during this period involve an initial engagement of the dying cell by Müller glia via the extension of processes from the Müller cell (Figure 1E), revealing that Müller glia are remarkably active in sensing and contacting dying cells.

### Apoptotic cargo and Müller glial cellular material is transferred from Müller glia to microglia

Our previous studies demonstrated that microglia dominate dying cell clearance in the developing zebrafish retina (Blume et al., 2020; Thiel et al., 2022b). Visualizing Müller glia in addition to microglia in real-time reveals that, although Müller glia primarily make the first contact (Figure 1), the process of dying cell engulfment often involves interaction of both Müller glia and microglia with the dying cells. In particular, we observed that SecA5-YFP+ cells initially engaged and enveloped by Müller cell processes are frequently commandeered by microglia (Movie 3, Movie 4; Figure 2). In a general synopsis of this sequence, the onset of YFP signal is met with contact by Müller cell processes forming a phagocytic cup-like structure that envelops the YFP+ target. During this process, migratory microglia are presumably in transit to the site of the YFP+ cell, and upon arrival, the microglial cell executes a maneuver by which the YFP+ cell enveloped by the Müller cell’s processes is transferred to the microglia cell for terminal engulfment (Figure 2A, 2B, Movies 3, 4).

**Figure 2:**
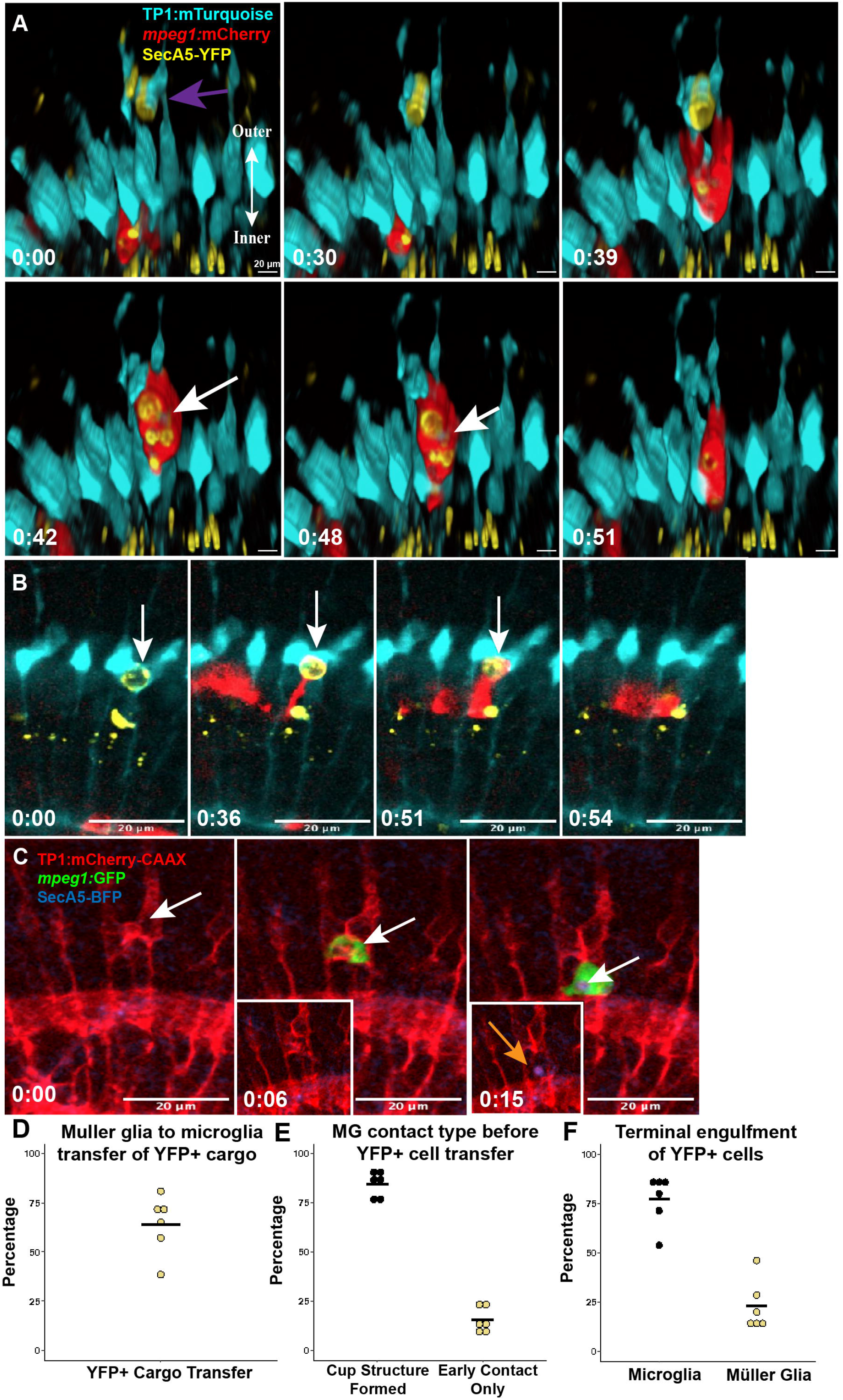
Transfer of cargo and cellular material from Müller glia to microglia. A: Time-lapse stills showing a Müller cell (cyan) engaging a YFP+ dying cell in the apical retina (purple arrow) and enwrapping the target with cellular processes and a cup structure before a microglial cell (red) emerges and retrieves the cargo. Pieces of the Müller cell cytoplasmic reporter are detected and move with the cargo (white arrow), suggesting cytoplasmic material transfer. B: Time-lapse stills show an inner retinal YFP+ cell being initially engaged by Müller glia, with a phagocytic cup-like structure formed around the cell before the YFP+ cell is transferred to microglia. C: Time-lapse stills showing microglial cell (green) interacting with a mCherry-CAAX+ (red) Müller glia compartment (white arrow). Microglia retrieves cargo from the Müller cell, taking mCherry membrane reporter (red) co-labeled with SecA5-BFP (orange arrow, inset, 0:15). C: Quantification of events in which YFP+ cargo is originally engaged by MG then commandeered by microglia, for each recording session, revealing that the majority of phagocytic events involve the transfer of dying cell cargo from Müller glia to microglia. Timestamps, bottom left (hr:min), relative to the first frame in the time series. D: The percentage events in which initial Müller cell contact with a YFP+ cell occurs prior to the transfer to and engulfment by microglia. E. Of events where Müller glia make the first target cell contact, the percentage of events in which this contact involves a cup structure versus a transient, more limited early contact. F: Terminal engulfment of SecA5-YFP+ cells by cell type.

Our quantifications show that the transfer of YFP+ cells initially contacted/enwrapped by Müller cell processes to microglia is a common occurrence in clearing dying cells from the normal, developing retina (Figure 2D). The majority of these YFP+ cell transfers occur after a cup-like structure has formed by Müller glia, though a smaller majority are engulfed by microglia after having only received early contact by Müller cell processes (Figure 2E). Ultimately, terminal engulfment is most commonly executed by microglia (Figure 2F), despite the initial contact being more frequently achieved by Müller glia (Figure 1C). As we observed this process of YFP+ engulfment and transfer, we also observed mTurquoise+ signal, representing the Müller cell cytoplasmic reporter, transferred to the microglia along with the YFP+ cargo (Figure 2A, white arrow). In other recordings, we again detected mTurquoise+ MG cytoplasmic reporter inside of microglial compartments (Movie 5, Figure 3A). Interestingly, this indicates that in addition to transfer of the enwrapped dying cells from Müller glia to microglia, some of the Müller cell cytoplasm is also transferred.

**Figure 3:**
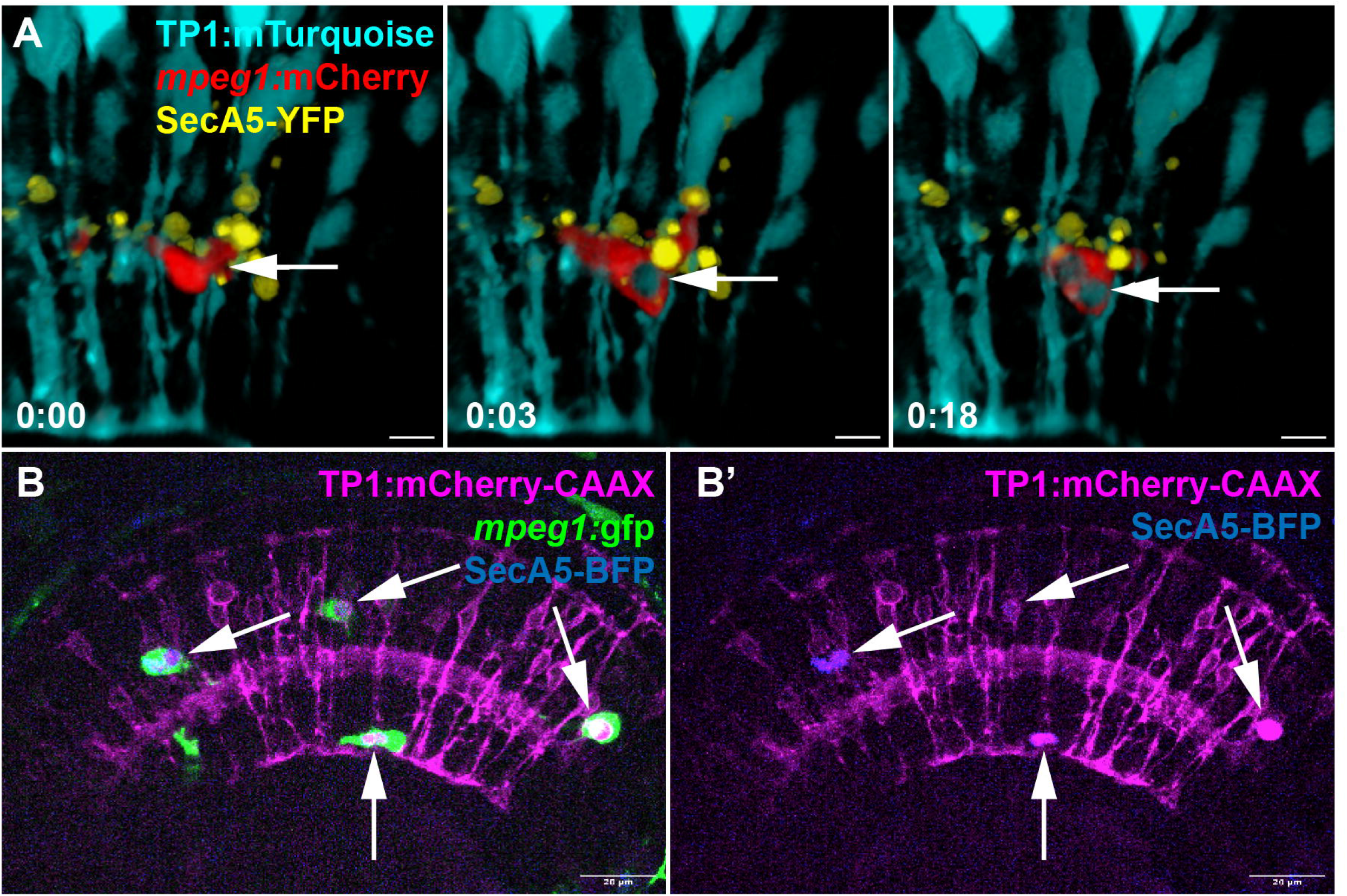
Detection of Müller glia components inside microglial compartments. A: In our recordings using *TP1:mTurquoise* as the Müller cell reporter, we also observed, at times, Müller cell cytoplasmic reporter inside of microglial compartments (arrow). Timestamps, bottom left (hr:min), relative to the first frame in the time series. Scale bar = 20 µm.B-B’: Using the membrane tagging reporter *TP1:mCherry*-CAAX, we observe Müller cell components colocalized with SecA5-BFP signal inside of microglial compartments. This signal is retained as microglia migrate through the retina.

To further visualize this interaction and cargo transfer, we used an alternative set of transgenic reporters in which the Müller glia were visualized by membrane-tagged mCherry (*TP1:mCherry-CAAX*), microglia with *mpeg1:gfp* (Ellett et al., 2011) and dying cells with *bact2:secA5-mTagBFP* (hereafter SecA5-BFP). In this triple transgenic system, the membrane tagged reporter demonstrated the formation of Müller cell process-derived compartments, which preceded the engagement by microglia and BFP signal (Figure 2B). Microglia were seen to engage with Müller cell compartments, with a concurrent detection of the BFP+ cell death reporter (Figure 2C, orange arrow). However, unlike the YFP reporter for PtdSer-exposing cells, the BFP reporter did not reliably report prior to compartment formation and the signal overall was weak, though it became more prominent once inside of microglia. We consider that differences in visualizing PtdSer-exposing cells with the YFP versus BFP reporters is a result of inherent differences in transgene drivers/expression, fluorophore brightness, as well as acid sensitivity (Subach et al., 2008; Young et al., 2010). Transfer of dying cell cargo from MG to microglia is supported as the onset of BFP signal occurs upon interaction of microglia with the MG compartment, and the BFP signal then travels with the microglial cell. Interestingly, what was most apparent from the combination of these reporters was the detection of Müller glia membrane-tagged mCherry signal within microglia compartments (Movie 6, Figure 3B. Strong mCherry+ membrane signal is persistently visible within the migratory microglia (Fig 2C, time 0:15, orange arrow). As stated above for the SecA5-YFP reporter, we again did not observe BFP+ Müller glia or collapse of Müller glia, indicating that the clearance of apoptotic Müller glia is unlikely to be a significant origin of the Müller cell reporter signals (cytoplasmic or membrane tagged) detected within microglia. Rather, the observations from both the MG cytoplasmic and membrane tagged reporters suggest that cellular material is transferred from MG to microglia concomitant with that of dying cell cargo.

### Müller glia complete engulfment of apoptotic cells in a subset of clearance events

The transfer of dying cell cargo from Müller glia to microglia, with the cargo terminally engulfed by microglia, is the predominant outcome in dying cell clearance (Figure 2C, 2E). Although less frequently observed, we did see in our recordings a subset of dying cells terminally engulfed and degraded by Müller glia (Figure 2E, Movie 7, Figure 4). In these instances, we observed a complete process of Müller cell extension towards, and around apoptotic cell bodies coupled with an inward retraction of the process that results in an internalization of the dying cell in a putative compartment, with compaction and fading of YFP signal inside of the MG cell body (Movie 7, Figure 4A).

**Figure 4:**
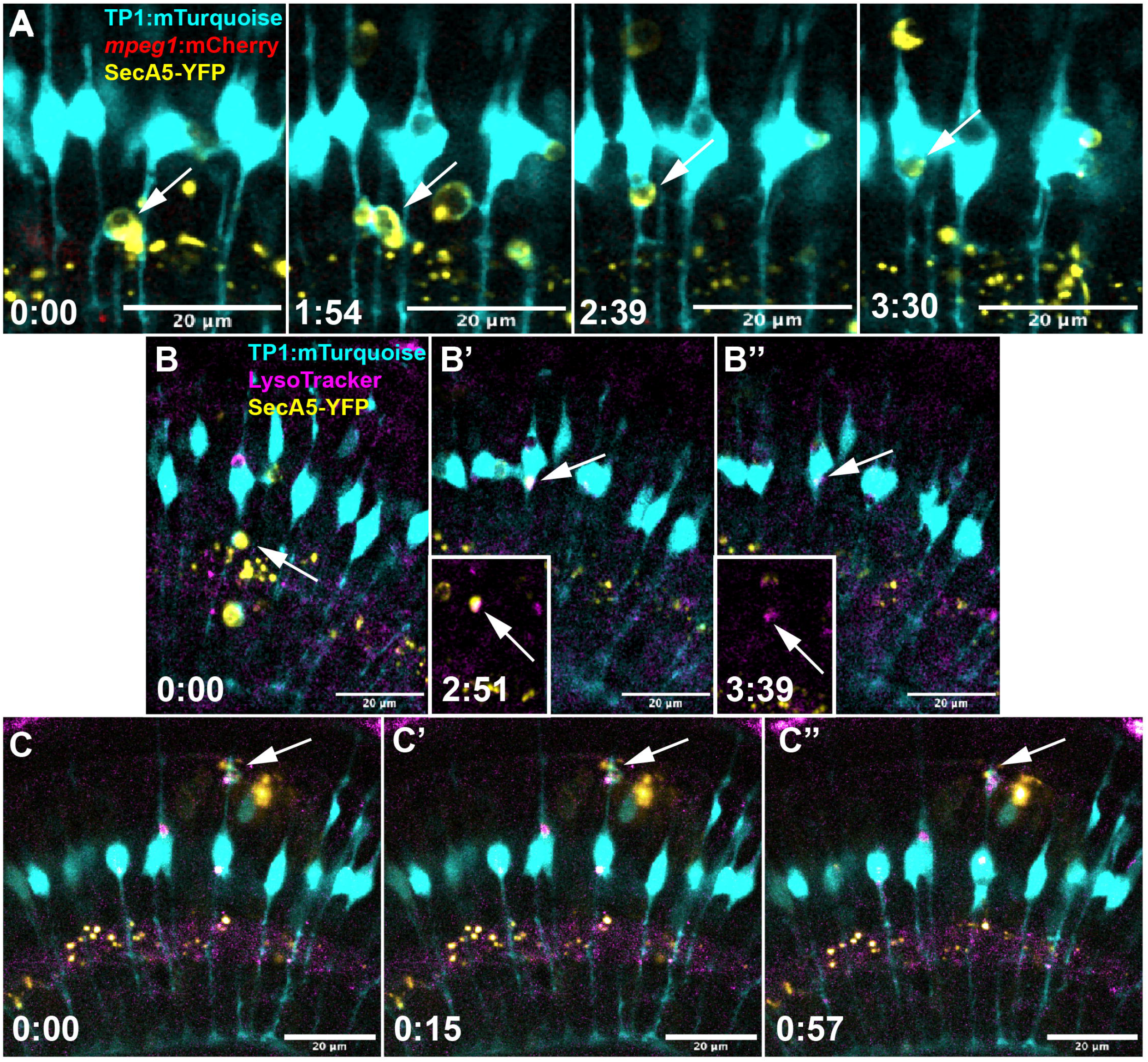
Müller glia complete engulfment of apoptotic cells. A: Time-lapse image stills show the formation of an mTurquoise+ phagocytic cup (cyan) around YFP+ dying cell (white arrow) followed by a progression of internalization into the Muller cell body. B-B”: Time-lapse stills show the process of Müller glia (cyan) engulfment of a YFP+ cell (white arrow). The YFP+ cell is brought into the Müller cell body upon which the compartment fluoresces with the lysosome signal (magenta, co-label in white). Insets show only YFP and magenta coloring to reveal co-localization of the YFP+ cell with lysosotracker (B’, inset) then loss of YFP+ signal (B’’, inset). C-C””: In some cases, YFP+ cells are engulfed and degraded by Müller glia (cyan) in a region other than the primary cell body; here shown to occur in the apical region of the Müller cell. Again, the YFP+ signal co-labels with the lysosomal reporter (magenta, indicated by white arrow). Timestamps, bottom left (hr:min), relative to the first frame in the time series.

To further confirm degradation of YFP+ targets within Müller glia via lysosomes, we incorporated LysoTracker dye into our live imaging protocol to identify the presence of acidic lysosomal compartments within the cell (Movies 8, S1, S2, Figure 4B and 4C). We found that LysoTracker co-localized with vacuolar compartments in Müller glia containing the engulfed YFP+ target cell signal, which is then degraded (Movies 8, S1, Figure 4 C-C”, D-D”). Collectively, we conclude that at least some Müller glia complete terminal engulfment degradation of engulfed dying cells, resolving controversy about such capacity, and supporting that Müller glia are indeed involved in final dying cell clearance in the developing retina.

### Terminal dying cell clearance in microglia versus Müller glia

Given that we observed both microglia and Müller glia completing terminal engulfment and degradation of YFP+ cells, we next compared the process by which these cell types complete clearance. We noted the time at which the YFP+ cell death reporter was first detected, defining this as the onset of the cell death signal. We then observed the YFP+ cell until it was terminally cleared by either a microglial or Müller glial cell, and we defined this as the point in which the YFP signal was displaced from its native site and fully internalized to the phagocyte. The time between the onset of the YFP signal to terminal internalization by the phagocytic cell is defined as the net clearance time for a dying cell. Apoptotic cells had a wide range of net clearance times whether they were cleared by Müller glia or microglia (Figure 5A), and this did not change significantly based on the cell type performing the terminal engulfment. However, net clearance time does not measure how fast the terminal phagocyte actually clears the cell once it has contacted it. Thus, we defined another subpoint within the time window of the net clearance time, which we defined as the formation and attachment of a phagocytic cup. From the contact by a cellular process, we measured the time elapsed before the YFP+ cell body had been displaced from its site and internalized to its respective terminal phagocyte, defining this timeframe as the engulfment time (Figure 5C and 5D). Of note, we measured time from the formation of the phagocytic cup by the glial cell that completed the terminal engulfment. When we quantified this process for each glial cell type (Figure 5B), we found a substantial difference between the speed of engulfment by microglia vs Müller glia, where microglia averaged ∼12 minutes to complete terminal engulfment, whereas Müller glia averaged ∼88 minutes (Figure 5B).

**Figure 5:**
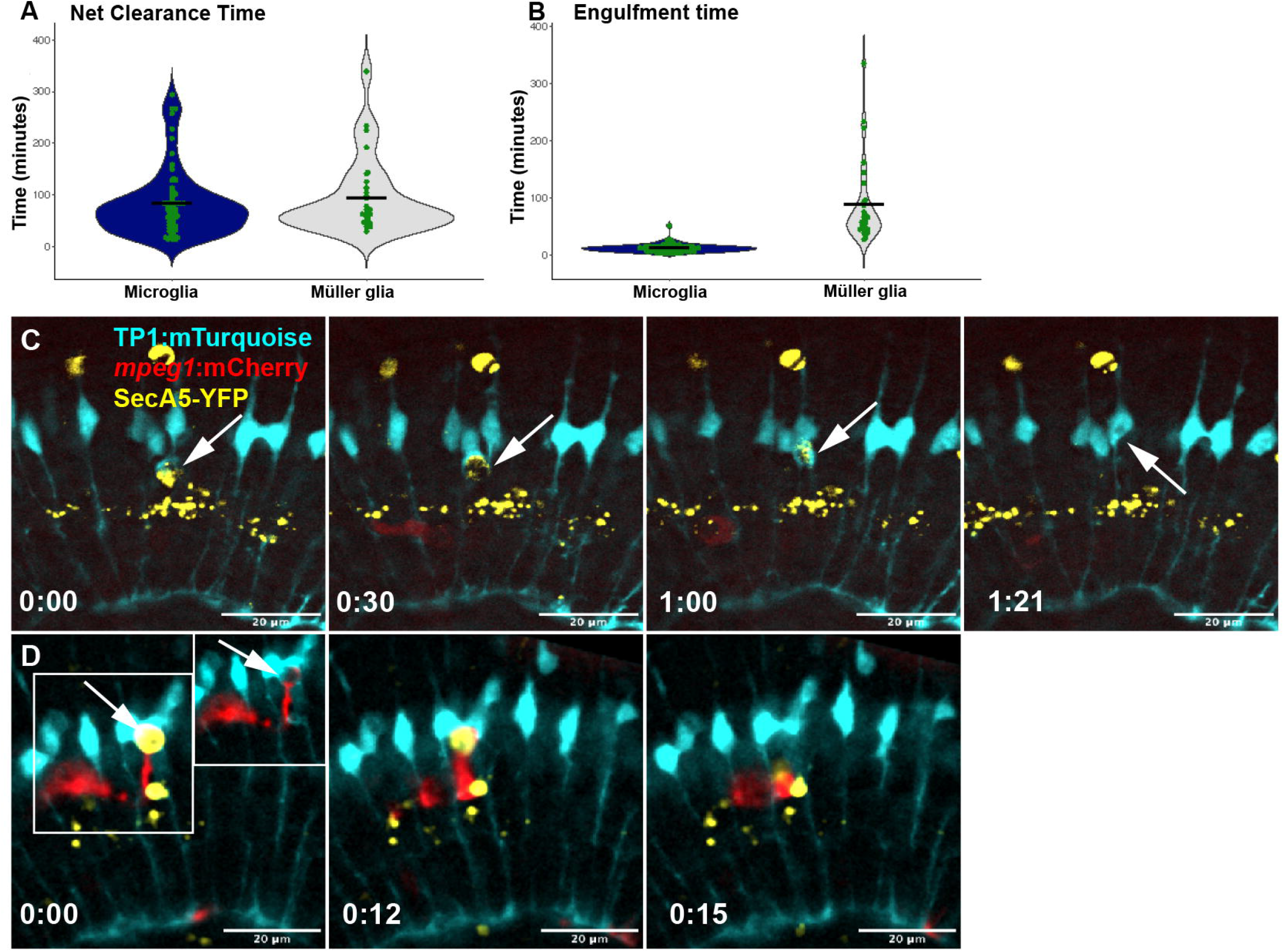
Quantification of clearance time and engulfment speed by cell type. A: Net clearance time: time from onset of apoptosis signal to terminal clearance by either microglia or Müller glia. B: Engulfment time: time from the attachment of phagocytic cup to the YFP+ target cell by either microglia or Müller glia, to engulfment into the glial cell. C and D. Examples of terminal engulfment by Muller glia (C) or microglia (D). C: Time lapse stills of Müller glia (cyan) terminal engulfment of a YFP+ cell, with time 0:00 coinciding to the formation of a phagocytic cup around the YFP+ cell. In this example, the Muller glial cell completes engulfment in 81 minutes. D: Time lapse stills of microglia (red) engulfment of a YFP+ cell, with 0:00 coinciding to the formation of a phagocytic cup around the YFP+ cell. In this example, the microglial cell completes engulfment in 15 minutes. Timestamps (C,D), bottom left (hr:min), relative to the first frame in the time series.

### Other Dynamic and Novel Müller Cell Behaviors

Our time-lapse recordings further revealed several dynamic, intriguing, and novel behaviors by Müller glia that to our knowledge have not previously been described. One such noted observation is the fragmentation or splitting of SecA5-YFP+ cargo coupled with distribution to other Müller glia (Movie 9, Figure 6A). In this instance, a single YFP+ target cell was moved to a single Müller cell body before it was fragmented and distributed among multiple adjacent Müller cells (Movie 9). Additionally, we observed instances in which at least two Müller glia reach for the same YFP+ target, resulting in an apparent “wrestling” or “tug-O’- war” between these two cells for the cargo (Figure 6B, Movie 10). Similarly, we may see that this wrestling may be coupled in the form of lateral passing of cargo between multiple Müller cells (Movie 11, Figure 6C). In addition to these behaviors of engulfment and cargo management, we observed the extension of remarkably long, dynamic, and winding cellular processes from the body of Müller cells (Movie 12, Figure 6D). These particular processes were visibly longer and more protrusive than the more commonly observed cellular process extensions used to engage proximal apoptotic cell bodies (described in Figure 1A, 1B, 1D).

**Figure 6:**
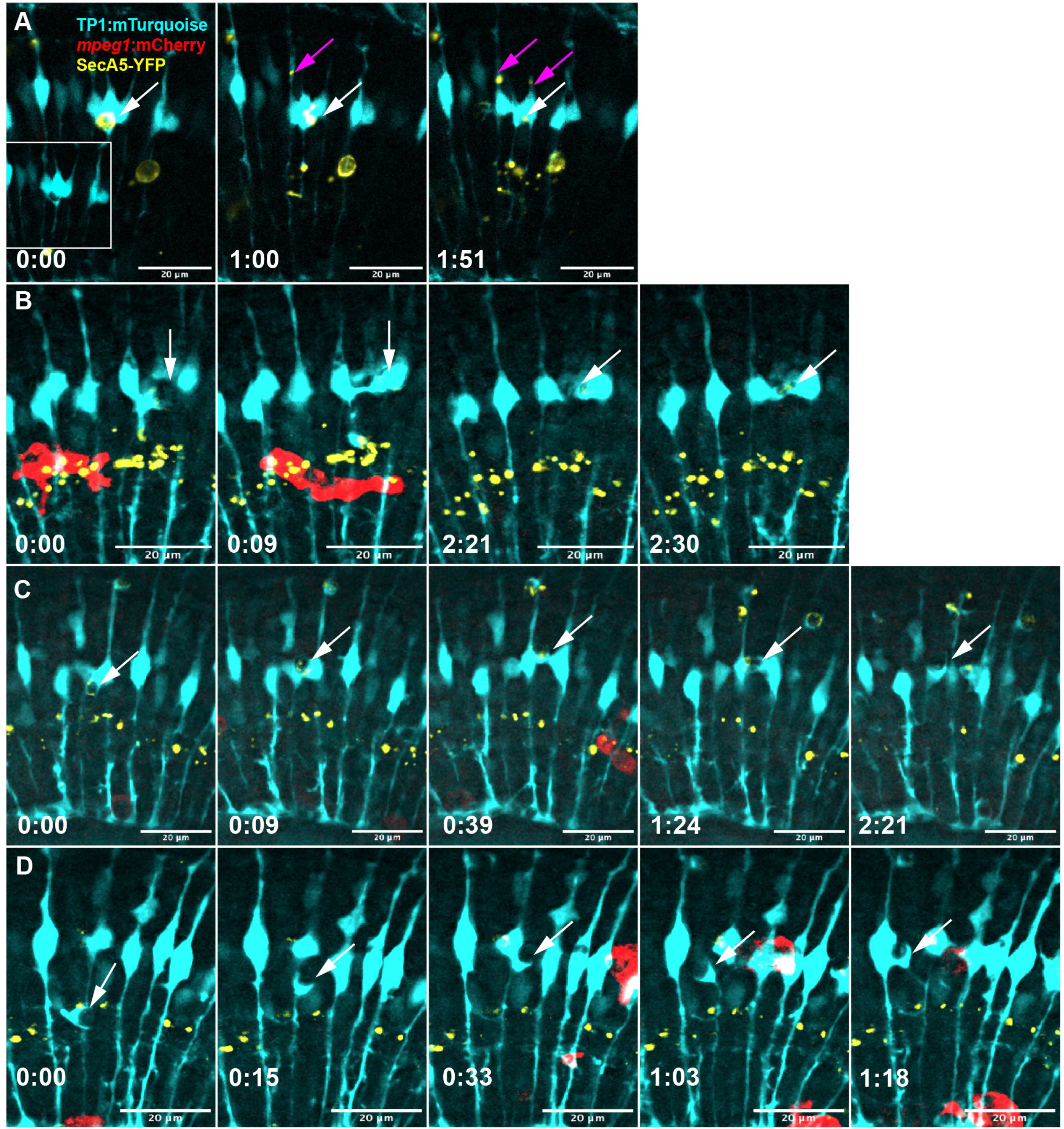
Other intriguing Müller glia behaviors revealed in real-time. A: Müller cell with a YFP+ cell (white arrow) fragments the cell and distributes the fragments to adjacent Müller cells (magenta arrows). B: Müller cell extends a tentacle process around an emerging YFP+ apoptotic cell and engages with another Müller cell before partitioning the cell (time 2:27 hr). C: An apoptotic cell is engaged by two Müller glia simultaneously. Over time the cell is captured by one of the initial two cells, but it is subsequently transferred to another cell after another bout of wrestling. D: A long extension protruding from a Müller cell (arrow) winds to and from adjacent cells back towards the original cell body over time. Timestamps, bottom left (hr:min), relative to the first frame in the time series.

### Clearance of phosphatidyl serine+ puncta in the inner retina

As described in our previous study (Blume et al., 2020), we observed numerous puncta (less than 3-micron diameter) from the SecA5-YFP reporter visualized in a location consistent with the inner plexiform layer of the developing retina (basal to the MG cell body yet apical to the MG end-feet, Figure 7). The presence of these SecA5-YFP+ puncta indicate that phosphatidyl serine (PtdSer)+ regions exist in the inner retina. Interestingly, localized exposure of PtdSer may occur at synapses that are pruned by microglia (Park et al., 2021; Rueda-Carrasco et al., 2023; Scott-Hewitt et al., 2020). These YFP+ puncta became visible independent of larger YFP+ cell bodies, and we observed that both microglia and Müller glia were active in the engulfment and clearance of the PtdSer+ puncta (Movie 13, and Figure 9). Müller glia engulf these SecA5-YFP+ puncta with fine process extensions and appear to internalize these puncta more efficiently compared to that seen for larger YFP+ cell bodies (Movie 13 (right), Figure 7B).

**Figure 7:**
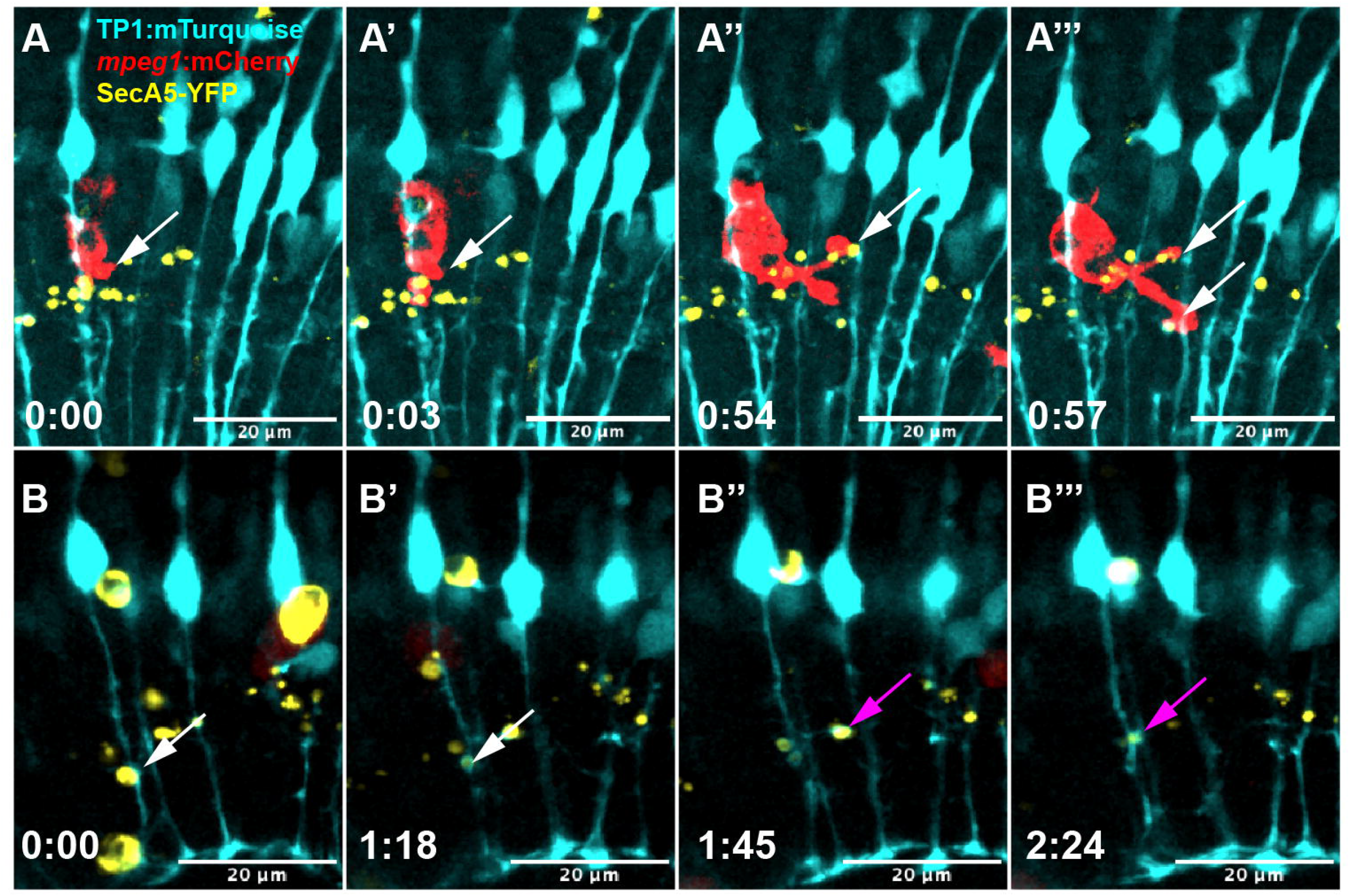
Microglia and Müller glia engulf PtdSer+ puncta in the inner plexiform region. A-A’’’: A microglial cell (red) engulfs distinct YFP+ puncta with multiple processes extended from the cell body (white arrows). B-B’’’: Müller glia (turquoise) also engulf YFP+ puncta using thin local processes. The white arrow shows one YFP+ puncta engulfed, then the same Müller cell engulfs a second YFP+ puncta (magenta arrow). Timestamps, bottom left (hr:min), relative to the first frame in the time series.

### Expression of selected phosphatidyl serine-recognizing receptors by microglia and Müller glia

Clearance of dying cells by phagocytes is achieved through receptor-ligand interactions between the phagocyte and apoptotic cell. Exposed phosphatidyl serine (PtdSer) on the surface of apoptotic cells serves as a ligand for PtdSer-binding phagocytic receptors on phagocytes. A wide array of phagocytic receptors exists and have, to varying extents, been characterized using various models or conditions of phagocytosis (Naeini et al., 2020). The existing body of work concerning microglial phagocytic receptors has largely been concentrated in mammalian models and have focused on the microglia of the brain (VanRyzin, 2021). Even in zebrafish studies, it has also been the brain rather than the retina that has been in focus (Mazaheri et al., 2014). In contrast, expression of phagocytic receptors by Müller glia is lesser known, though recognition of PtdSer is likely involved (Nomura-Komoike et al., 2020; Thiel et al., 2022b). In addition, phagocytic receptors expressed by zebrafish retinal microglia has not been directly examined.

We therefore examined expression of phagocytic receptors by microglia and Müller glia during zebrafish retinal development, to suggest those which could be important for phagocytic behaviors observed in our recordings.

We used hybridization chain reaction fluorescent in situ hybridization (HCR-FISH, (Choi et al., 2018)) to probe expression of mRNA encoding selected phagocytic receptors known to bind to PtdSer either directly or through bridging molecules (Naeini et al., 2020). These receptors were selected based on expression in RNA-seq datasets probing zebrafish microglia and Müller glia (Mitchell et al., 2019; Oosterhof et al., 2017; Sifuentes et al., 2016) and experimental reports (Anderson et al., 2022; Hsiao et al., 2019; Mazaheri et al., 2014; Nomura-Komoike et al., 2020). The selected transcripts encode *axl, mertka, havcr1, timd4, itgam.1, itgb2,* and *lrp1aa*. The products of the genes *mertka* and *axl* belong to a family of receptor tyrosine kinase proteins known as the TAM family (Lemke, 2017). These proteins bind to PtdSer via the bridging molecule Gas6 (Lemke, 2017). The zebrafish genes *havcr1* and *timd4* belong to a group of proteins known as T-cell immunoglobulin domain-containing proteins (TIM) and bind PtdSer directly (Naeini et al., 2020; Xu et al., 2016); these genes have orthology to mammalian TIM4 and TIM1 (Xu et al., 2016). *Itgam.1* and *itgb2* are predicted orthologs of human/mouse ITGAM and ITB2, which together in mammals form the Complement Receptor 3 (CR3) (Bader et al., 2021). Binding of CR3 is bridged with PtdSer via C1q(Naeini et al., 2020). LRP1 can act as a phagocytic receptor (Donnelly et al., 2006) and was upregulated by Müller glia responding to retinal damage (Sifuentes et al., 2016).

We visualized mRNA in situ in the developing zebrafish retina at 3 days post fertilization, coinciding with the endpoint of our timelapse recordings, in combination with microglia (*mpeg1*:mCherry and/or *mpeg1* transcript) and Müller glia *(TP1*:mTurquoise) specific markers. We found that microglia expressed multiple PtdSer-recognizing receptors, with strong expression of *axl, havcr1, mertka, and itgb2* (Figure 8A, B, C, D). Interestingly, while expression of *itgb2* was strong and detectable in all microglia, the expression of *itgam.1* was only detected in a subset of microglia and even in those cells, the signal was sparse and dim (Figure 8E). It is possible that in zebrafish, the complement system does not directly mirror that seen in mammals and may employ a different integrin-alpha binding partner for *itgb2*, or *itgb2* is used differently. In contrast to that seen for microglia, we found that Müller glia expressed only two of these selected PtdSer receptors (*mertka* and *itgb2,* Figure 8C, D’), this expression was apparently heterogenous, and the expression did not match the level found in microglia. We did not detect appreciable expression of *timd4* in any cells in the retina at this timepoint (Figure 8F), while expression of *lrp1aa* appeared nearly ubiquitous in the retina (Figure 8F) and was not restricted to any particular cell type, glia or other. Collectively, these results indicate redundant and strong expression of multiple PtdSer receptors by microglia, with more limited expression by Müller glia. Such expression patterns are consistent with the dominant role of microglia in terminal engulfment and could underlie the basis for cargo transfer from Müller glia to microglia or to other MG more competent for engulfment.

**Figure 8:**
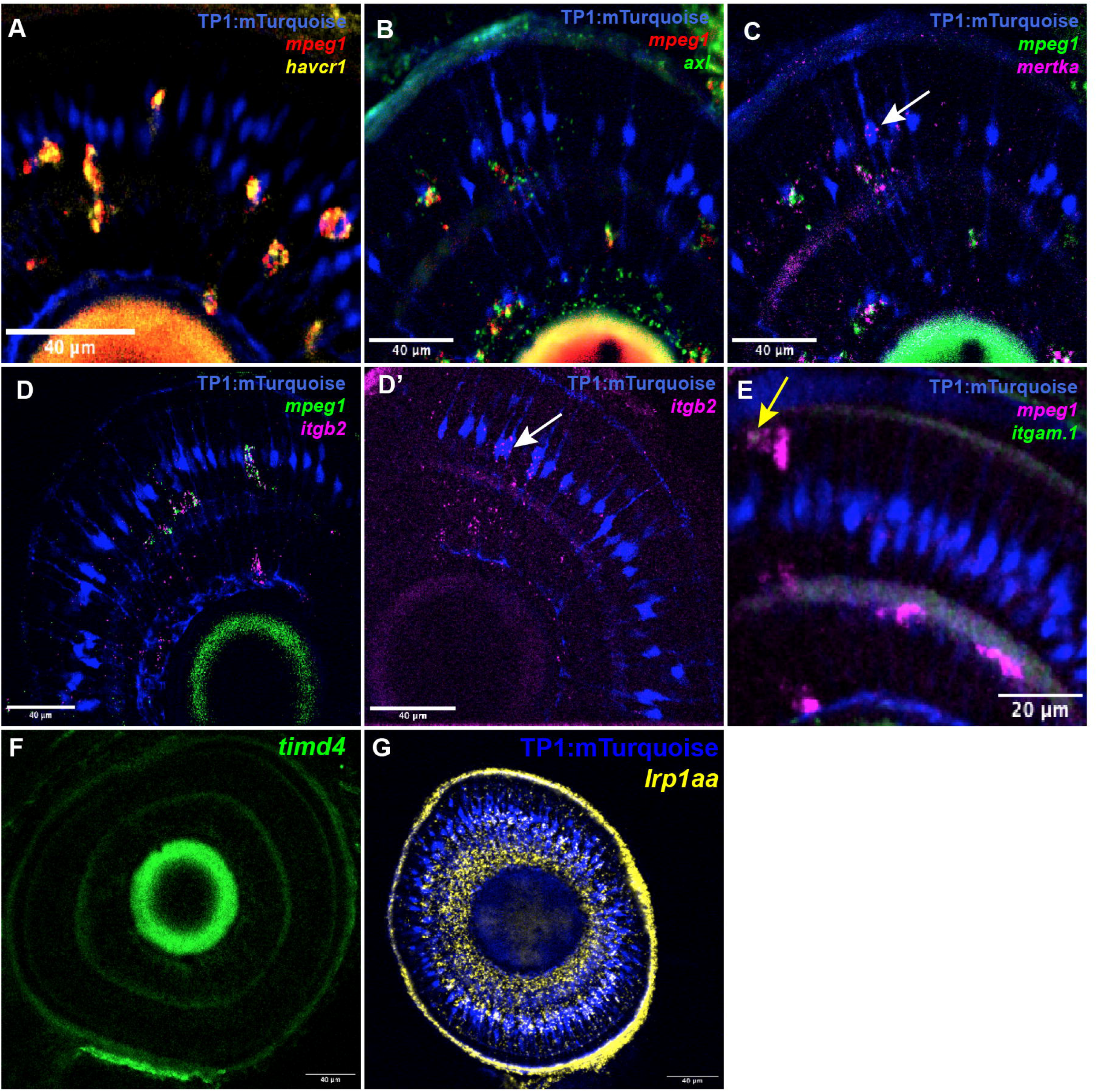
Expression of selected phagocytic receptors in wild-type zebrafish retinas during development. HCR-FISH was used to visualize transcripts for the indicated receptors in retinas expressing markers for Muller glia (*TP1:mTurquoise*, blue) and microglia (*mpeg1*:mCherry/*mpeg:1* transcript) at 3 dpf. Images show selected z planes acquired from whole embryonic eyes/retinas. A: Expression of *havcr1* was limited to microglia and highly expressed. B: The receptor *axl* was also strongly expressed in microglia cells. C: Expression of *mertka* was found in both microglia and Müller glia, though microglia displayed stronger expression, and expression in Müller glia expression appeared heterogeneous, with clusters of Müller showing transcript label in the cell body. D-D’: Complement receptor component *itgb2* was strongly expressed in microglia (D) in addition to some expression by Müller glia (D’), albeit in heterogeneously and in varying amounts. E: Prospective complement receptor pair member *itgam.1* was faintly expressed in subset of microglia, and weakly so (arrow). F-G: Expression of *timd4* (F) and *lrp1aa* (G).

## Discussion

Prior to this work, simultaneous visualization of Müller glia, microglia, and dying cells in real-time in the intact developing retina has not been described. Though we have examined microglial clearance of dying cells (Blume et al., 2020) and others have recorded microglial clearance of ablated rod photoreceptors (White et al., 2017) in developing retinas, real-time imaging of the Müller glia is limited to their developmental differentiation (MacDonald et al., 2015). Our current study further exploits the advantages of the zebrafish for live timelapse imaging to observe behaviors of Müller glia and microglia together in response to programmed cell death occurring during retinal development. We describe through our recordings the phagocytosis and clearance of these dying cells, illustrating a complex and novel set of interactions between Müller glia and microglia. The results of our study reinforce the paradigm of microglial dominance in phagocytosis; however, we additionally reveal that Müller glia are surprisingly dynamic and active in responding to dying cells and observed that both glial cell types interact with most apoptotic cells prior to their clearance. Not only do the Müller glia most frequently make the first contact with the apoptotic cell, they also frequently form an initial phagocytic cup-like structure around the target, after which the target is then often engulfed and terminally degraded by microglia following a transfer process. Despite the frequent “switch” of phagocyte for terminal engulfment after initial contact, Müller glia do indeed fully execute phagocytosis for a subset of dying cells. Collectively, these real-time observations resolve seemingly conflicting reports from fixed tissue by revealing transient yet common interactions of both glial cell types with a target cell and resolving that both cell types do indeed complete phagocytic engulfment.

In between the beginning and end of the apoptosis-phagocytosis sequence, an average of over 60% of all phagocytic events involved “switch” of phagocyte from Müller glia to microglia. It is not known whether this is a coordinated or unilateral process. At times microglia were engaging and commandeering the apoptotic cell before the Müller glia could completely enwrap the cell, yet more commonly, there was a formed cup-like structure around the dying cell at the time of microglial engulfment. Furthermore, the detection of Müller glial cell reporters, both cytoplasmic and membrane-tagged, being taken from Müller glia to microglia with the apoptotic cell raises new questions over the structure of the compartments formed, and potential biological basis for such transfer, given that we did not observe apoptotic Müller glia in our recordings. In particular, such observations suggest that the entire Müller cell cup structure may be engulfed by microglia in addition to the apoptotic cell target. If material transfer occurs and is a biologically important process remains to be determined. However, it is worth noting that intercellular transfer from photoreceptors to Müller glia (Hutto et al., 2023), and from macrophages to other cells (Kidwell et al., 2023; Roh-Johnson et al., 2017), have been documented. Nonetheless, the observations from our recordings support that both a “strong-armed” approach by microglia, in which microglia outcompete Müller glia for the apoptotic cell to pull it away, may be possible in addition to a coordinated process in which one or both cells execute a regulated signaling process for the exchange of the YFP+ cells, both of which are yet to be investigated.

Our findings indicate that sensing of dying cells and their phagocytosis, when executed, is an active process on the part of Müller glia rather than passive collapse around already enwrapped neurons. Müller glia actively extended cellular processes towards phosphatidyl-serine (PtdSer) exposing cells, often times prior to detection of the cell death reporter on the targets. The molecular sensing of targets by Müller glia remains unknown but may follow suit with regards to apoptosis-sensing mechanisms found in macrophage cells including, but not limited to, purinergic sensing (Blume et al., 2020), fractalkine (Sokolowski et al., 2014), and lysophosphatidylcholine (Lauber et al., 2003). After contact, phagocytic cups frequently formed, and in scenarios when the Müller cell completed engulfment, this was followed by a subsequent retraction inward. Our usage of a lysosome tracker in real-time confirmed the active degradation of target cells by Müller glia upon lysosomal fusion. This is consistent with our previous work showing Rab5+ endocytic vesicles and increased lysosomal staining inside of Müller glia in fixed tissue samples from microglia-deficient retinas, where Müller glia show compensatory phagocytic activity (Thiel et al., 2022b). In the mouse brain, several reports indicate the capacity of astroglia to perform phagocytosis of apoptotic cells and their debris (Damisah et al., 2020; Konishi et al., 2020; Konishi et al., 2022; Morizawa et al., 2017). Given that studies in the invertebrate *C. elegans*, which lack professional phagocytes like macrophages, served as a foundation for our knowledge of genes regulating the molecular process of dying cell clearance (Conradt et al., 2016; Lukácsi et al., 2021), it is not entirely surprising that vertebrate neuroglia display such capacity. It is suggested that vertebrates developed specialization for managing phagocytic load through the use of professional phagocytes such as macrophages/microglia, possibly to reduce load on neuroglia, to in turn allow more specialized and complex neuroglial functions in the vertebrate central nervous system. Such an idea is consistent with signs of reactivity from Müller glia due to increased phagocytic load when microglia are absent (Thiel et al., 2022b).

In the case of microglial phagocytosis of apoptotic cells, we see a single microglial cell manage an entire cellular target in terms of acquisition and degradation. On the other hand, Müller glia exhibited apparent varied behaviors, strategies, or mechanisms for handling phagocytic loads. In some cases, engulfment was not completed by a sole Müller glia, but rather by a consortium that involved more than one Müller cell either wrestling for the cell, or partitioning the cell in a way in which more than one cell received a portion to ingest. In some cases, Müller cells apparently passed the targets to neighboring Müller cells. Yet, in other cases, a single Müller cell would complete the engulfment of the cell. In this light, it could be said that microglial phagocytosis is more uniform in its procession than Müller glial phagocytosis. In contrast, for PtdSer+ puncta, Müller glia appeared to efficiently engulf these smaller targets.

This may suggest that heterogeneity in phagocytic capacity exists for Müller cells, and this variable capacity for engulfment could exist at a cellular level and depend on the size of the target.

In addition to the relative phagocytic loads, there were other factors that separated the phagocytic characteristics of microglia and Müller glia. One such factor is the relative time of engulfment. Microglia were noticeably faster in completing the clearance of apoptotic cell bodies upon the attachment of a phagocytic cup. This process took significantly longer for Müller glia. This difference may be owed to the greater ability of microglia to adapt their morphology for process extension and migration via rapid cytoskeletal rearrangements, whereas Müller glia are more or less conformed to a fixed radial shape, with more steps required for process extension and phagocytic cup formation. We did observe many times that Müller glia are capable of extending dynamic processes from various parts of their cell body. An additional difference could derive from the expression of PtdSer-recognizing receptors. Of the selected receptors investigated, microglia strongly express at least four of them (*axl, mertka, itgb2, havcr1)*, while Müller glia were limited to the expression of *merkta* and *itgb2.* Yet even for those two genes, expression strength did not visibly match that of microglial cells and expression amongst Müller glia was not apparently uniform. Heterogeneity in *mertka* and *itgb2* expression by Müller glia is also represented in publicly available single cell-RNAseq datasets (Hoang et al., 2020) and could explain apparent differences in phagocytic capacity between Müller cells. Further, the difference in expression of PtdSer receptors between the two glial cell types may play a role in determining the nature of the phagocytic dominance of microglia and these receptors may play a potential role in cargo switching between the two cell types in which microglia remove the target from the Müller glial phagocytic cup.

Another interesting observation, though not the intended focus, was the clearance of PtdSer+ puncta in an area that is consistent with the inner plexiform layer, a laminated region of synaptic connections in the inner retina. While our studies do not explicitly identify the nature of this puncta, existing literature suggests that localized PtdSer exists on synapses and could be recognized by microglia for synaptic refinement (Györffy et al., 2018; Kurematsu et al., 2022; Park et al., 2021; Rueda-Carrasco et al., 2023; Scott-Hewitt et al., 2020). In our recordings, PtdSer+ puncta were engulfed by both microglia and Müller glia. If these puncta do indeed visualize synapses, then our recordings suggest that both microglia and Müller glia could be involved in synaptic clearance during retinal development, at least in zebrafish. In the brain, astroglia have been observed to directly mediate the synaptic refinement through phagocytosis (Chung et al., 2013; Yang et al., 2016), though shared roles between multiple glial cell types have still not been well examined.

Collectively, our recordings reveal tremendous insight into previously unknown behaviors of the Müller glia, some of which suggest functional roles that have not yet been explored. Such dynamic and interactive behaviors should change our view of how the Müller glia function in the vertebrate retina, and better inform our understanding of these glial cells in health and disease.

## Materials & Methods

### Animals

All procedures were in compliance with protocols approved by the Institutional Animal Care and Use Committee at the University of Idaho. Adult zebrafish were housed in an aquatic environment at 28.5°C with monitored, recirculating system water and an automated diurnal light cycle of 14 hours of light and 10 hours of darkness according to (Westerfield, 2007).

Zebrafish embryos were collected from breeder pairs with spawning time corresponding to the onset of light in the housing environment. Embryos were collected and housed in an incubator at 28°C with a final concentration of 0.06 % N-Phenylthiourea (PTU) added to the water to prevent the development of melanin pigment. Water and PTU solutions were refreshed daily. At an age of at least 2 days post-fertilization (dpf), embryos were sorted for positive expression of each transgene; embryos expressing all desired transgenic reporters were selected for use in the experiments. Prior to usage for experiments, the chorion was manually removed with fine tip forceps as necessary. Zebrafish cannot be sexed until reproductive age, therefore sex of embryos is unknown.

### Transgenic Lines

The *TP1:mTurquoise* (uoi2523Tg) transgenic line used to visualize Müller glia was generated in-house using a Tol2 transgenesis construct kindly gifted by Ryan MacDonald (University College London) based on (MacDonald et al., 2015). The *TP1:mCherry-CAAX* transgenic line was described in our previous work (Thiel et al., 2022b).Microglia were visualized using the pre-existing transgenic lines *mpeg1*:mCherry (gl23) or *mpeg1*:GFP (gl22) (Ellett et al., 2011), originally obtained from Zebrafish International Resource Center (ZIRC) and maintained at the University of Idaho. Apoptotic cells were visualized using the TBP:Gal4;UAS:SecA5-YFP transgenic line (referred to throughout this paper as SecA5-YFP) (van Ham et al., 2010) originally gifted by Randall Peterson (University of Utah) and maintained at the University of Idaho. We generated the transgenic line *bact2*:SecA5-mTagBFP (uoi2518Tg) using Gateway cloning to recombine p5E-*bact2* (obtained from the 2007 Tol2 kit) (Kwan et al., 2007), pME-secAnxaV-NS (Morsch et al., 2015), Addgene), and p3E-mTagBFP(Don et al., 2017), Addgene) into pDEST-Tol2CG2 (Tol2 kit). The size and orientation of the final construct was validated by restriction enzyme digestions. For in-house generated transgenic lines, single blastomere-stage zebrafish embryos were microinjected with ∼0.5-1 nL of DNA construct mixed with Tol2 mRNA (25 ng/µl final concentration). F0 founders were selected based on expression of the transgene as embryos, grown to adults, then out-crossed with non-transgenic wildtype fish to F1 showing germline inheritance of the transgene. F1 were then again out-crossed to obtain stable lines at F2 or later generations, with ∼50% transgene segregation upon out-cross to non-transgenic partners. Zebrafish carrying multiple transgenic reporters were obtained by crossing stable transgenic lines over two to three generations.

### Live-time lapse imaging

For timelapse recordings, embryos were anesthetized in a solution of water, 0.06% PTU, and 0.016% Tricaine (MS-222). Once anesthetized, embryos were suspended in liquified, low-melting temperature agarose (1% agarose prepared in fish system water) and overlayed onto a coverslip-bottom dish (No. 1.0 thickness, 35mm, MatTek Corporation). Embryos were oriented so that they were on their side with one eye against the coverslip. The agarose gel pad, which then solidified, was overlayed with water/PTU/MS-222 solution to ensure continued anesthesia and pigment inhibition of the fish during imaging.

Embryos on the coverslip dishes were transferred to a climate-controlled chamber (Okolab) maintained at 28°C on the microscope stage. Images were acquired using a Nikon Andor spinning disk confocal microscope connected with a BSI Express 16-bit sCMOS camera. Imaging was performed using a CFI APO LWD 40x water immersion 1.15 NA λ S DIC N2 objective. An immersion compound (Zeiss Immersol™ W) with a refractory index of 1.334 was placed on the objective in lieu of water to avoid evaporation over the imaging session. The region for imaging was selected based upon the visibility of entire radial Müller glia. Imaging depth spanned approximately 40µm, with 3 µm z-steps. Each imaging session lasted 10 hours with 3-minute time intervals between each capture, spanning a total of 12-15 z-stacks. Embryos were selected for imaging and analysis by display of strong heartbeat and circulation throughout the tissue both before and after the imaging session. The mTurquoise, YFP, and mCherry fluorophores were excited with 445 nm, 514 nm, and 561 nm lasers, respectively, with a Chroma(TM) 89006 (eCFP, eYFP, mCherry) dichroic mirror. For imaging GFP, mCherry, and BFP, excitation used 488 nm, 546 nm, and 405 nm, respectively and the Chroma (TM) 89000 Sedatquad dichroic mirror.

### Image viewing and processing

Timelapse image stacks were viewed with both Nikon Elements software and FIJI. Images selected for figures were derived from selected time points in a recording and converted from their original 16-bit format into a RGB tiff format after image adjustments using FIJI. 3D renderings for selected movies were performed using Nikon Elements through projection in volume view. Movies were made by saving select time series as “.avi” files with jpeg compression. Images were assembled into figure arrangement and annotated in Adobe Illustrator. Movies were annotated using FIJI.

### Quantification of phagocytic events in timelapse recordings

Images were quantified manually using ImageJ software. Counts were made by noting phagocytic events during the recording by examining individual z-stacks and maximum z-projection images to avoid overcounting. Phagocytic events for apoptotic cells were defined as the terminal engulfment of SecA5-YFP+ bodies (of size 4-7 microns in diameter) by a glial cell. Each event was then studied in terms of the glial cell that contacted the cell first (by watching the image series prior to the terminal engulfment) and the glial cell that completed terminal engulfment of the YFP+ cell. Each cell that was engulfed during a recording session was then scored for first cell type to contact it (Müller glia or microglia), visual evidence of transfer from one cell type to the other, and the final cell type to terminally engulf it (Müller glia or microglia). The transfer of cargo was defined as YFP+ cells that were initially engaged by Müller glia but were subsequently engulfed by microglia.

The net clearance time was recorded as the elapsed time beginning at the timepoint at which a YFP+ cell appeared in the recording until the timepoint at which it was terminally cleared by a glial cell. This elapsed time and the phagocyte cell type (Müller glia or microglia) was recorded for each YFP+ cell analyzed. YFP+ cells already present at the beginning of the recording, or which were not terminally cleared by the end of the recording, were not analyzed because true elapsed clearance time could not be determined. The engulfment time for each glial cell (Müller glia or microglia) was determined as the elapsed time beginning at the time the glial cell formed a phagocytic cup around the YFP+ cell to the timepoint at which it successfully removed the apoptotic cell from its native site and internalized it. The engulfment time elapsed and the terminal phagocyte cell type (Müller glia or microglia) was recorded for each YFP+ cell analyzed. Cells already being internalized by phagocytes prior to the beginning of the recording, or those not completed by the end of the recording, were not counted in the analysis. Graphical presentation of quantifications was performed using RStudio.

### Lysotracker™ Immersion

To visualize lysosomes in live zebrafish retinas, we prepared embryos the same as described above with an additional step prior to mounting. In this step, a 1µM solution of LysoTracker™ Deep Red (made from 1 mM stock in zebrafish system water) was prepared with 0.06% PTU. Embryos were placed in this lysotracker/PTU solution and incubated in the dark at 28°C for 30 minutes. Following incubation, embryos were washed twice with system water to remove excess LysoTracker™ from the surface of the embryos. Embryos were then placed in a new solution of system water and 0.06% PTU prior to suspension in the agarose pad. Embryos were oriented in the agarose pad and mounted on the coverslip dish as described above. The agarose pad containing the embryos was overlayed with 1µM LysoTracker™ plus 0.06% PTU in fish system water. For imaging of LysoTracker™ Deep Red, a 640 nm (far-red) excitation laser was used.

### Fluorescence In-Situ Hybridization

RNA fluorescent in-situ hybridization was performed using hybridization chain reaction (HCR) protocol, reagents, and probes from Molecular Instruments (Los Angeles, USA) (Choi et al., 2018) and as performed previously to examine microglia-expressed transcripts in zebrafish embryos (Thiel et al., 2022a). Probes were custom designed by Molecular Instruments based upon each transcript’s NCBI accession number (Table 1). At 3 dpf, embryos were fixed in a freshly made solution of 4% paraformaldehyde (PFA) in 1x phosphate buffered saline (PBS). Prior to fixation, the PFA/PBS solution was cooled to 4°C to prevent autofluorescence in the samples. Once in the fixative solution, the embryos were incubated for 24 hours at 4°C with constant rocking. After fixation, the PFA solution was removed, and the embryos were washed three times with RNAse-free 1x PBS solution for 5 minutes per wash. Samples with transgenic fluorescent reporters (e.g., *TP1:mTurquoise*) were protected from exposure to light to prevent destruction of the fluorophore. Following washes, embryos were washed 4 times with 100% methanol for 10 minutes each, followed by one 50-minute wash in 100% methanol. Embryos were stored in a fresh solution of 100% methanol and stored at -20°C at least overnight prior to use.

**Table 1.** NCBI accession number, probe set size, and hairpins used in HCR in situ hybridization.

| Transcript | NCBI Accession Number | Probe set size | Amplifier format/fluorophore |
| --- | --- | --- | --- |
| *mpeg1* | NM_212737.1 | 17 | B1/546, 488; B3/546,488 |
| *mertka* | XM_002664231.6 | 20 | B2/647 |
| *axl* | XM_017351109.2 | 20 | B3/AF488 |
| *havcr1* | NM_001002434.1 | 18 | B3 AF546 |
| *itgb2* | XM_680920.7 | 20 | B2/647 |
| *itgam.1* | XM_009306407.4,<br>XM_009306408.3,<br>XM_005156007.4,<br>XM_009306409.3 | 17 | B1/488 |
|  | (probes will<br>detect all) |  |  |
| *timd4* | NM_001122617.1 | 19 | B3/AF488 |
| *lrp1aa* | XM_021479679.1 | 20 | B1/AF546 |

To begin the hybridization procedure, embryos were rehydrated in a graded series of methanol: PBS-Tween (1x PBS + 0.1% Tween 20, PBST) washes for five minutes each. The order of the graded series (in ratio of MeOH: PBST) follows: 75:25, 50:50, 25:75, and finally 100% PBST. Following the rehydration of the embryos, they were treated with 1 ml of a 10 µg/ml solution of proteinase K for 10 minutes at 37°C. Proteinase K solution was removed, and embryos were rinsed briefly with PBST twice at room temperature. This was followed by post-fixation with 4% PFA in PBS for 20 minutes at room temperature. Embryos were then washed five times with 1x PBST for five minutes each. PBST was replaced by probe hybridization buffer that had been pre-heated to 37°C, and embryos were pre-hybridized for 30 minutes at 37°C. A solution of probes was made using 500 µl of probe hybridization buffer and 2 pmol of each probe, with appropriate combinations for multiplexing. After pre-hybridization, the solution was removed, and embryos were incubated with the probes in hybridization solution for 12-16 hours at a temperature of 37°C in a hybridization oven.

Following the hybridization, probe solutions were removed, and embryos were washed with probe wash solution four times for 15 minutes each at 37°C, followed by two washes with 5x SSCT (saline sodium citrate-0.1% Tween-20) buffer for five minutes each. Amplification was performed by incubation in probe amplification buffer for 30 minutes at room temperature, followed by an overnight incubation (>12 hours) with amplifier solution. 30 pmol of each hairpin (see Table 1) was snap cooled by incubation at 95°C for 90s in a thermocycler then cooled to room temperature in the dark for 30 minutes. Snap-cooled amplifiers were added to amplification buffer at room temperature as appropriate for multiplexing. The pre-amplification buffer was replaced with the hairpin mixture and incubated for 12-16 hours at room temperature, protected from light. Hairpins were removed, and embryos were washed with SSCT five times, with DAPI added at 1:1000 in the final wash, then stored in 1x PBS at 4°C until imaging.

The embryos/eyes were subsequently mounted either with or without dissection of the eyes from the body and suspended in glycerol on a 22×60 mm microscope coverslip, using two smaller 22.5×22.5 mm coverslips to act as spacers, then covered and sealed with another 22×60 mm coverslip. Samples were imaged using either 20x air objective or the 40x water-immersion objective lens (described above) on the Nikon Andor spinning disk confocal microscope described above with 2µm z-stacks. Images were collected using Nikon Elements and subsequently processed using ImageJ.

## Acknowledgments

Funding for this work was provided by NIH (R01 EY030467, D.M.M). We thank Dr. Onesmo Balemba (Director of the IDAC Imaging Core, University of Idaho) and the Institute for Interdisciplinary Data Sciences/Initiative (IIDS/IBEST) for support of our research. We thank current and former members of the Mitchell lab for technical support and care of zebrafish, and Whitney Thiel for blastomere microinjection. We thank Dr. Ryan MacDonald (University College London) for gifting the *TP1:mTurquoise* and *TP1:mCherry-CAAX* transgenesis constructs, and Dr. Deb Stenkamp (University of Idaho) for draft review of our paper.

